# IscR-mediated morphological regulation confers virulence and stress resistance by reducing stress molecule uptake in *Acinetobacter baumannii*

**DOI:** 10.64898/2026.05.04.722593

**Authors:** Hoan Van Ngo, Nayoung Kim, Jumi Park, Jinki Yeom

**Affiliations:** Department of Biomedical Science, College of Medicine, Seoul National University, Seoul, Korea; Department of Microbiology and Immunology, College of Medicine, Seoul National University, Seoul, Korea; Cancer Research Institute, Seoul National University, Seoul, Republic of Korea

## Abstract

Living organisms must adequately respond to stress to survive and proliferate. Bacterial pathogens face multiple stressors during infections, including oxidative stress from host innate immune cells and antibiotic treatment from clinical therapy. The pathogenic bacterium *Acinetobacter baumannii* is considered the most critical threat to public health due to its broad antibiotic resistance. However, it is poorly known how *A. baumannii* properly responds to antibiotics and stress molecules during infection. Here, we investigate the mechanisms by which *A. baumannii* regulates its morphology to reduce the uptake of stress molecules under oxidative stress and antibiotic exposure, thereby conferring virulence and survival during infection. The transcriptional regulator IscR responds to oxidative stress by upregulating *pbp1a*, which encodes an enzyme involved in peptidoglycan biosynthesis. Under oxidative stress, bacteria undergo a morphological shift from a rod to a coccoid form, reducing their surface area and thus decreasing their absorption of reactive oxygen species. Inactivation of either *iscR* or *pbp1a* results in an elongated morphology characterized by an elevated surface area, thereby reducing *A. baumannii* survival under oxidative stress. Furthermore, IscR-mediated morphological control is essential for survival under antibiotic treatment. Moreover, IscR-mediated morphology regulation is required for *A. baumannii* survival in macrophage and mouse models. These findings elucidate a strategy by which *A. baumannii* uses IscR to adapt to stress through morphological control, facilitating its survival during infections against both immune response and antibiotic therapy.

**IMPORTNACE:** *Acinetobacter baumannii* is a major cause of nosocomial infections. It poses a critical threat due to its extensive antibiotic resistance. This study reveals that the pathogen can change its cellular shape to survive immune system attacks and antibiotic treatment. This change represents a previously unknown survival strategy. *A. baumannii* transitions to a coccoid morphology under oxidative stress and antibiotic treatment. It does so by activating the peptidoglycan synthesis gene *pbp1a* through the IscR transcriptional regulator. This rapid morphological adaptation helps *A. baumannii* evade host defenses and resist antibiotic treatment by reducing uptake of stress molecules. Our findings advance understanding of how pathogens adapt to hostile environments and identify new therapeutic targets. By blocking this shape remodeling ability, it may be possible to render pathogenic bacteria more vulnerable to immune responses and antimicrobial treatments. This offers a promising strategy for combating this multidrug-resistant pathogen.

## INTRODUCTION

All living cells meticulously change their shape in response to environmental conditions (1–5). For instance, they detect ambient osmolarity and subsequently alter their cellular shape by regulating transcription, translation, metabolism, and membrane dynamics (1–5). Bacteria can alter their cellular shape to resist stressful environments by regulating elongation, division, and cell wall synthesis pathways (1–5). During infections, pathogenic bacteria encounter harsh conditions in the host, including oxidative stress induced by reactive oxygen species (ROS) produced by innate immune cells and antibiotic treatment during clinical therapy (6–11). Thus, regulation of cellular shape in response to ROS and antibiotics is essential for bacterial survival during infections (12–15). Nonetheless, it remains largely unexplored how pathogens modulate their morphology in response to various stress environments during infection. In this study, we show that pathogenic bacteria rapidly alter cellular morphology in response to stress conditions to enhance survival under antibiotic exposure and during host immune responses.

Bacteria can alter their morphology to endure stress and adapt to environmental changes (1–5). The bacterial cell wall, composed of peptidoglycan, is crucial for preserving cell shape and viability (16). In proteobacteria, the elongasome and divisome are two multiprotein complexes that coordinate peptidoglycan production and assembly, thereby regulating bacterial cell length and the development of daughter cells (17, 18). The class A penicillin-binding proteins (aPBPs), PBP1a and PBP1b, are bifunctional enzymes with both glycosyltransferase (GTase) and dd-transpeptidase (dd-TPase) activities, which play a significant role in the synthesis of the cell wall and septal peptidoglycan (16). In *E. coli*, PBP1a and PBP1b exhibit functional redundancy in regulating cell morphology, as the deletion of either protein does not affect bacterial morphology (19, 20). By contrast, in *A. baumannii*, which poses a particular risk to immunocompromised patients and is associated with hospital outbreaks due to its high rate of antibiotic resistance (21–25), PBP1b is unable to substitute for the function of PBP1a (26). *A. baumannii* PBP1a collaborates with PBP3 to synthesize septal peptidoglycan, which is crucial for cell division and the formation of daughter cells (26). Although the biochemical functions of aPBPs in peptidoglycan synthesis in *A. baumannii* have been characterized, their physiological role in the bacterial stress response remains unclear.

In bacteria, the transcriptional regulator IscR is an essential component of the *isc* operon for the biogenesis of iron-sulfur (Fe-S) clusters (27, 28). IscR utilizes Fe-S clusters as cofactors to respond to oxidative stress (29–32). Under normal conditions, holo-IscR, which contains the Fe-S cluster, represses the *isc* operon (27, 28). The holo-IscR binds to both Type 1 and Type 2 DNA motifs in promoter regions of target genes (30). In contrast, oxidative stress damages the Fe-S cluster in IscR. This damage leads to the production of apo-IscR. Apo-IscR activates the *isc* operon and initiates Fe-S cluster biogenesis, restoring cellular homeostasis (27, 32, 33). Apo-IscR exclusively binds to Type 2 DNA motifs in target promoter regions (28, 33–35). Moreover, apo-IscR regulates not only the *isc* operon but also activates additional systems in response to oxidative stress. For instance, apo-IscR directly activates the *sufA* operon, an alternative Fe-S cluster biogenesis system in *E. coli* (33), and the hemolysin synthase *vvhBA* operon in *Vibrio vulnificus* (36). Thus, IscR is a crucial regulator of stress responses and virulence in diverse bacterial pathogens (32, 35–38). Nevertheless, the pathogenic role of IscR in various pathogens, including *A. baumannii,* remains unexamined, despite the bacterium’s significance in antibiotic resistance and clinical settings.

This study demonstrates that *A. baumannii* IscR directly activates the *pbp1a* gene expression in response to oxidative stress and antibiotic treatment, enabling the bacteria to change a coccoid morphology. Coccoid morphology reduces the uptake of stress molecules, which enhances survival under oxidative stress and antibiotic exposure. Inactivation of *iscR* disrupts coccoid morphology, leading to the emergence of filamentous bacteria that are more vulnerable to oxidative stress and antibiotics. IscR directly binds to the *pbp1a* promoter region, promoting a morphological transition that reduces surface area and stressor uptake under stress conditions. IscR-mediated morphological regulation is essential for bacterial survival within macrophages and in the murine infection model. Together, our findings reveal a novel regulatory mechanism in which pathogenic bacteria transcriptionally modulate their cellular morphology to reduce the uptake of stress molecules, thereby enhancing their survival and proliferation during host infections.

## RESULT

### IscR is required for defense against oxidative stress and virulence during infection by regulating cellular morphology

All domains of life require an iron-sulfur assembly system to generate iron-sulfur (Fe-S) clusters, which serve as critical cofactors for various enzymes (39–42). Reactive oxygen species (ROS) can damage iron-sulfur clusters, leading to their disruption and subsequent loss of iron from enzymes. Bacteria have an Fe-S repair system, represented by the *isc* operon, to maintain the function of various enzymes (41, 43). *Acinetobacter baumannii*, a pathogenic bacterium, presents a high risk to immunocompromised patients due to its high rate of antibiotic resistance (44). The host immune system and antibiotic treatment cause oxidative stress (6–11). Given this, *A. baumannii* must overcome oxidative stress to cause disease in the host. Thus, we hypothesized that IscR, the master transcriptional regulator of the *isc* operon (Fig. 1A), is essential for the oxidative stress response and virulence during infections in *A. baumannii*.

**Figure 1.**
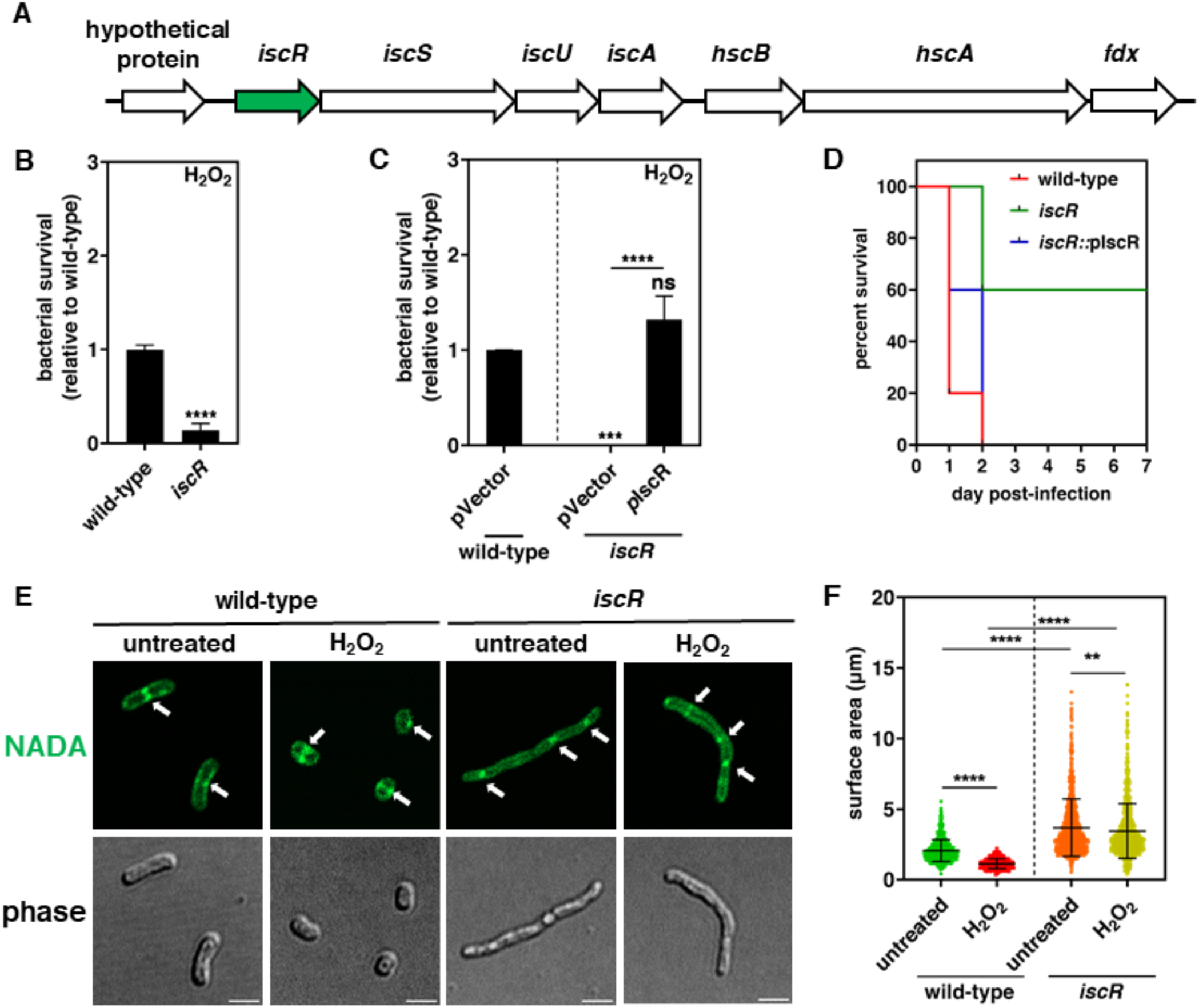
IscR-mediated morphological change of *A. baumannii* in response to oxidative stress. (A) The schematic image shows the *isc* operon in *A. baummannii*. The transcriptional regulator *iscR* gene (green color) is the first gene in the operon. (B) Relative survival of *A. baumannii* 17978 wild-type and *iscR* after treatment with 10 mM H_2_O_2_ for 2 h. Data are presented as means ± standard deviation (SD) from n = 3. *P* values were determined using an unpaired Student’s t-test. \*\*\*\**p* < 0.0001. (C) Relative survival of *A. baumannii* 17978 wild-type harboring the empty plasmid vector (pVector) and *iscR* harboring the empty plasmid vector (pVector) or the *iscR*-expressing plasmid (pIscR) after treatment with 10 mM H_2_O_2_ for 2 h. Data are presented as means ± SD from n = 3. *P* values were determined using one-way ANOVA with Tukey’s multiple comparison test. \*\*\**P* < 0.001, \*\*\*\**P* < 0.0001, ns; not significant. (D) Survival curves of mice infected with *A. baumannii* 17978 wild-type harboring the empty plasmid vector (pVector) and *iscR* harboring the empty plasmid vector (pVector) or the *iscR*-expressing plasmid (pIscR). Six-week-old BALB/c mice (n = 6) were challenged intraperitoneally with 5×10^8^ CFU/mL of bacteria, and the survival time was monitored. (E) Fluorescence (upper) and phase (lower) microscopy of *A. baumannii* 17978 wild-type and *iscR* in normal condition (untreated) or treated with 10 mM H_2_O_2_ for 30 min. NADA-green is integrated into the peptidoglycan of living bacteria during cell wall synthesis (45). Arrows indicate bacterial septa. Scale bar = 2 µm. (F) Surface area quantifications of each cell population (n ≥ 500) were calculated using ImageJ software with the MicrobeJ plugin. Each dot represents one bacterial cell. *P* values were determined using one-way ANOVA with Tukey’s multiple comparison test. \*\*\*\**P* < 0.0001, ns; not significant.

Multiple lines of evidence indicate that IscR is essential for oxidative stress defense and virulence in *A. baumannii*. First, the inactivation of *iscR* resulted in reduced bacterial survival under hydrogen peroxide (H_2_O_2_) exposure, known to be the cause of oxidative stress, in comparison to the wild-type (Fig. 1B). Second, the survival defect of the *iscR* mutant in H_2_O_2_ was restored by a plasmid expressing *iscR* through a heterologous promoter, whereas the empty plasmid vector did not restore survival (Fig. 1C). Third, mice intraperitoneally inoculated with wild-type showed survival decrease to 0% by day 2 (Fig. 1D, red line. By contrast, *iscR* inactivation attenuated bacterial virulence, resulting in 60% survival in mice at day 7 post-infection (Fig. 1D, green line). The *iscR* mutant strain harboring a plasmid expressing *iscR* via a heterologous promoter fully restored the virulence of *A. baumannii* (Fig. 1D, blue line). Collectively, these findings indicate that IscR is required for the survival under oxidative stress and virulence during infection in *A. baumannii*.

Then, we investigated how IscR contributes to oxidative stress defense and virulence in *A. baumannii*. Intriguingly, *A. baumannii* exhibited a transformation in cellular morphology from a rod to a coccoid form in response to oxidative stress (Fig. 1E) that reduces bacterial surface area (Fig. 1F). In the presence of H_2_O_2_, rod-shaped cells transitioned to a coccoid morphology, as demonstrated by peptidoglycan staining with the fluorescent D-alanine derivative NADA [NBD-(linezolid-7-nitrobenz-2-oxa-1,3-diazol-4-yl)-amino-D-alanine] (NADA-green) (Fig. 1E). NADA-green is integrated into the peptidoglycan of living bacteria during cell wall synthesis (45), indicating that the observed coccoid bacteria were viable. By contrast, the *iscR* mutant exhibited an elongated filamentous morphology with/without oxidative stress. In addition, the *iscR* mutant failed to transition to a coccoid form under oxidative stress conditions (Fig. 1E) and reduce bacterial surface area (Fig. 1F). The filamentous morphology of the *iscR* mutant is more sensitive to oxidative stress than the coccoid form in wild-type (Fig. 1B). Together, *A. baumannii* changes its morphology to coccoid to reduce its surface area via IscR, which enhances bacterial survival under oxidative stress and virulence during infection.

### IscR directly activates *pbp1a* expression to regulate cellular morphology

Next, to investigate how IscR controls cellular morphology in *A. baumannii*, we focused on the class A penicillin-binding protein 1a (PBP1a), which is essential for division and fitness in *A. baumannii* (26, 46). *A. baumannii* requires the PBP1a enzyme to preserve rod-shaped morphology (26, 46). This requirement depends on PBP1a peptidoglycan synthase activity at the septum. However, it is unclear how *A. baumannii* controls the *pbp1a* gene expression in response to stress. We found that the transcriptional regulator IscR is needed to regulate morphology under oxidative stress (Fig. 1). Consistent with this, we reasoned that IscR controls *pbp1a* expression during oxidative stress.

Multiple lines of evidence support the notion that IscR directly activates transcription of the *pbp1a* gene. First, inactivation of either *iscR* or *pbp1a* results in filamentous morphology (Fig. 2A) and increased cellular surface area (Fig. 2B), consistent with previous research on the role of PBP1a in the division of rod-shaped *A. baumannii* (26, 46). In contrast, the inactivation of *pbp1b*, which is a different penicillin-binding protein, did not affect the cellular morphology and surface area in *A. baumannii* (Fig. 2A and B). Also, peptidoglycan staining with NADA-green revealed multiseptated filamentous shape in *iscR* and *pbp1a* mutants, unlike the wild-type and *pbp1b* mutants (Fig. 2A) (26, 46). Second, a putative type II IscR binding site was detected upstream of the predicted transcriptional start site of *pbp1a* (Fig. 2C), which resembles the *E. coli* IscR type II motif (33, 34, 47). Third, luciferase reporter assays indicated that *pbp1a* expression was significantly increased under oxidative stress conditions (Fig. 2D). In addition, *pbp1a* expression dramatically decreased in the *iscR* mutant compared to the wild-type strain in both normal and oxidative stress conditions (Fig. 2D). By contrast, the expression of the *isc* operon is significantly elevated in the *iscR* mutant strain due to derepression under normal conditions, as opposed to oxidative stress conditions (Fig. 2E), which is consistent with previous findings (27). Fourth, the purified IscR protein binds to a DNA fragment harboring the promoter region of *pbp1a* (Fig. 2F). Furthermore, excess unlabeled *pbp1a* promoter DNA successfully competed out for binding with the labeled *pbp1a* promoter (Fig. 2F). In contrast, as expected, purified IscR was found to bind to the promoter region of the *isc* operon (Pisc) (27), but it did not bind to the promoter region of the *hscB-hscA-fdx* operon (P*hscB-hscA-fdx*), which does not have putative IscR binding site on the promoter region (Fig. 2F). Lastely, both *iscR* and *pbp1*mutants demonstrated increased minimal inhibitory concentrations (MICs) to all *β*-lactam antibiotics tested, except meropenem, compared with the wild-type and the *pbp1b* mutant (Table 1). Most *β*-lactam antibiotics target the dd-TPase activity of PBP1a, except meropenem, which targets both dd-TPases and ld-TPases (46). Thus, *iscR* and *pbp1* mutants cannot produce the target protein, PBP1a, which inhibits bacterial survival to *β*-lactam antibiotics. In contrast, the MIC of the membrane-targeting antibiotic colistin remained unchanged (Table 1). Taken together, IscR directly binds the *pbp1a* promoter and is required for *pbp1a* expression, which is essential for regulating cellular morphology in response to oxidative stress.

**Figure 2.**
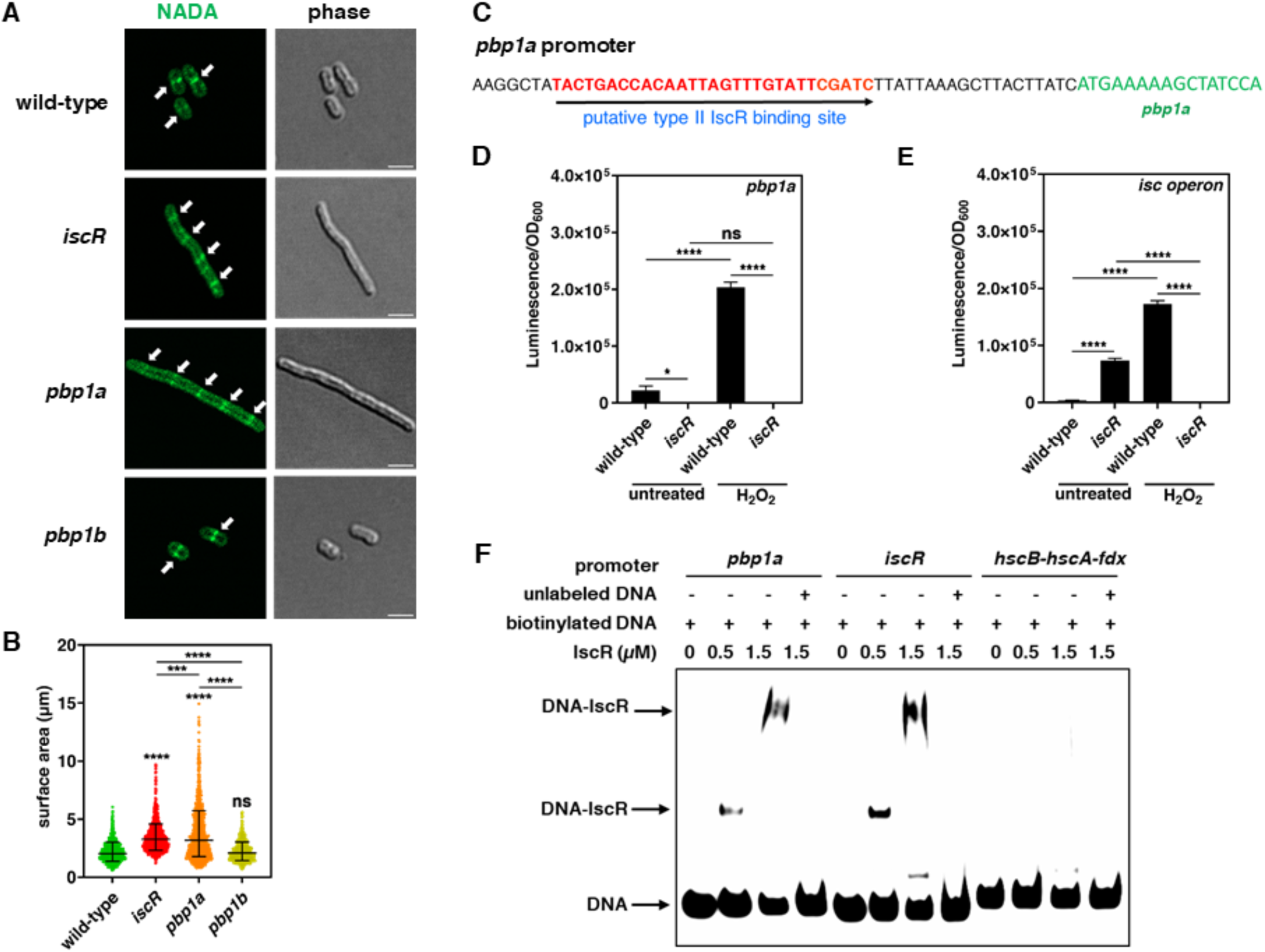
IscR directly activates *pbp1a* expression in *A. baumannii*. (A) Fluorescence (left) and phase (right) microscopy of *A. baumannii* 17978 wild-type, *iscR*, *pbp1a*, and *pbp1b* in normal conditions. Arrows indicate bacterial septa. Scale bar = 2 µm. (B) Surface area quantifications of each cell population (n ≥ 500) were calculated using ImageJ software with the MicrobeJ plugin. Each dot represents one bacterial cell. *P* values were determined using one-way ANOVA with Tukey’s multiple comparison test. \*\*\*\**P* < 0.0001, ns; not significant. (C) Putative IscR binding site on the promoter region of *pbp1a*. (D-E) Luminescence emission of *A. baumannii* 17978 carrying the luciferase reporter plasmid pLPV1Z fused with *pbp1a* promoter (D) or *isc* operon’s promoter (E) at 2 h in the absence or presence of 10 mM H_2_O_2_. Data are presented as means ± SD from n = 3. *P* values were determined using two-way ANOVA with Tukey’s multiple comparison test. \*\*\*\**P* < 0.0001, ns; not significant. (F) EMSA validates that IscR interacts with the promoters of *pbp1a*and *isc* operons, but not *hscB-hscA-fdx* operon. Representative images from three independent experiments with similar results are shown.

**Table 1.** Antibiotic susceptibilities of *A. baumannii* 17978 and its mutants.

| antibiotics |  |  | MIC (mg/liter) for strain: |  |  |  |
| --- | --- | --- | --- | --- | --- | --- |
| class | name | target | wild-type | *iscR* | *pbp1a* | *pbp1b* |
| B-lactams | carbenicillin | dd-TPase | 16 | 32 | 32 | 16 |
|  | cefoperazone | dd-TPase | 128 | 1024 | 512 | 128 |
|  | cefotaxime | dd-TPase | 32 | 64 | 64 | 32 |
|  | mecillinam | dd-TPase | 64 | 256 | 128 | 64 |
|  | meropenem | dd-Tpase and ld-TPase | 2 | 2 | 2 | 1 |
| lipopeptide | colistin | lipid A | 2 | 2 | 2 | 2 |

### PBP1a-mediated coccoid transition reduces ROS uptake that enhances bacterial survival

Here, we determined that *A. baumannii* undergoes a morphological transition from rod to coccoid shape in response to oxidative stress via IscR-mediated PBP1a expression, thereby improving bacterial survival under these conditions (Fig. 1-2). Since ROS molecules, such as H_2_O_2_, could diffuse across bacterial membranes (48), we hypothesized that reduced bacterial surface area, as seen in a coccoid shape, via PBP1a, may decrease the uptake of oxidative stress molecules to increase tolerance to stress and bacterial survival.

To investigate this hypothesis, we constructed bacterial strains harboring a plasmid expressing *pbp1a* via a heterologous promoter. The coccoid cell shape and reduced cellular surface were observed in strains harboring a plasmid expressing *pbp1a* under a heterologous promoter, without stress (Fig. 3A and B), similar to the coccoid morphology induced by ROS (Fig. 1E and F). Notably, the coccoid shape of bacteria significantly reduces the uptake of ROS, as indicated by measurements using intracellular ROS indicator, 2’,7’-dichlorodihydrofluorescein diacetate (DCFH-DA) (Fig. 3C), thereby enhancing bacterial survival (Fig. 3D). Subsequently, we treated sulbactam, which a class A *β*-lactamase inhibitor that targets both PBP1 and PBP3 in *A. baumannii* (49), to induce filamentous shape (Fig. 3E and F) like *pbp1a* mutant strain (Fig. 2A). These cells exhibited significantly increased ROS uptake (Fig. 3G) and demonstrated dramatically increased susceptibility to oxidative stress (Fig. 3H). Together, PBP1a-mediated morphology transition to a coccoid shape reduced the uptake of stress molecules that increase *A. baumannii* survival under oxidative stress.

**Figure 3.**
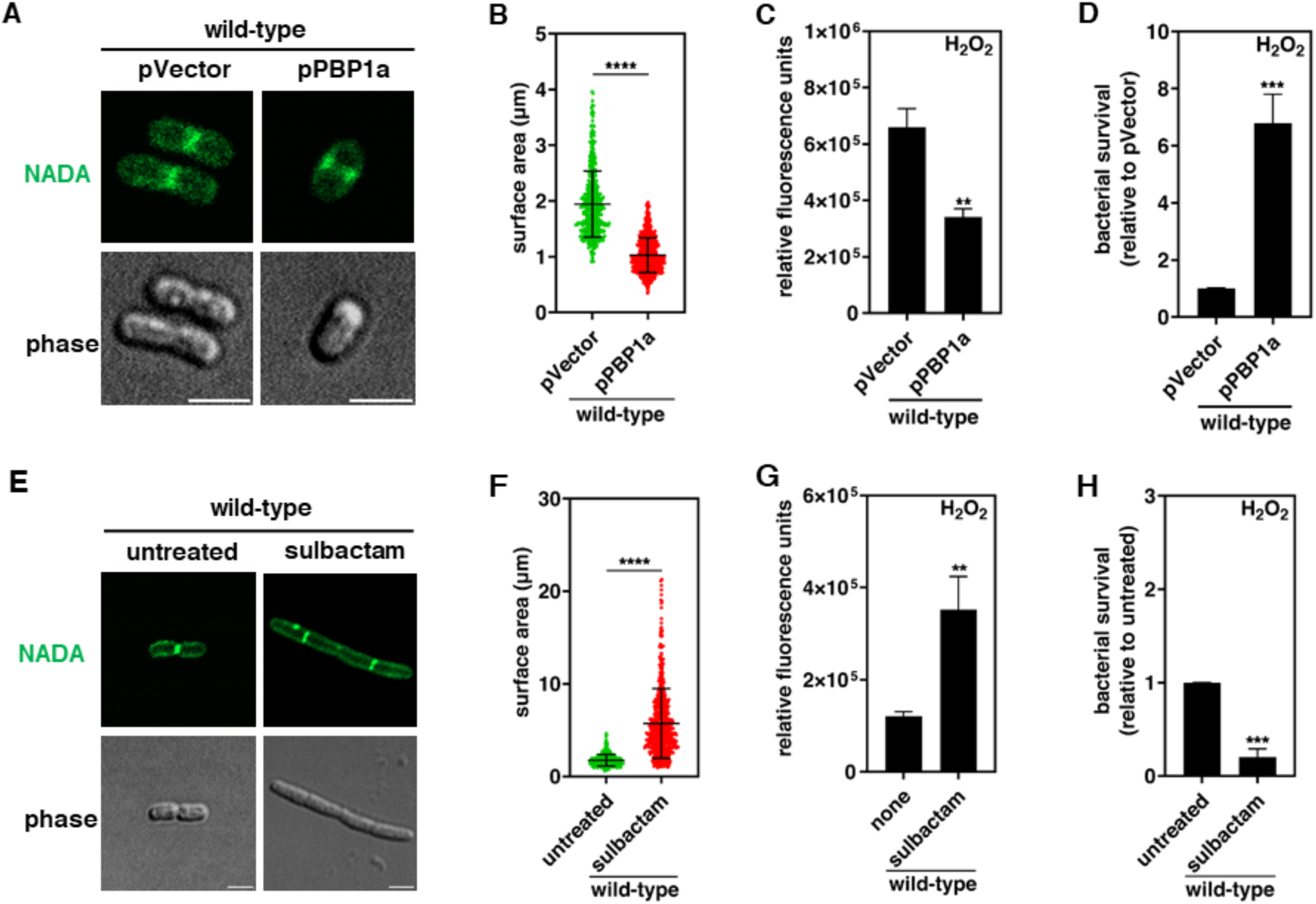
Coccoid morphology is required for bacterial survival under oxidative stress in *A. baumannii*. (A) Fluorescence (upper) and phase (lower) microscopy of *A. baumannii* 17978 wild-type harboring the empty plasmid vector (pVector) or the *pbp1a*-expressing plasmid (pPBP1a) in normal conditions. NADA-green is integrated into the peptidoglycan of living bacteria during cell wall synthesis (45). Scale bar = 2 µm. (B) Surface area quantifications of each cell population (n ≥ 500) were calculated using ImageJ software with the MicrobeJ plugin. Each dot represents one cell. *P* values were determined using an unpaired Student’s t-test. ****p < 0.0001. (C) ROS uptake of *A. baumannii* 17978 wild-type harboring the empty plasmid vector (pVector) or the *pbp1a*-expressing plasmid (pPBP1a), as measured by fluorescence intensity of DCF-DA upon treatment with 10mM H_2_O_2_ for 30 min. The results are expressed as relative fluorescence units, with the RFU value for the untreated control subtracted. Data are presented as means ± SD from n = 3. *P* values were determined using an unpaired Student’s t-test. \*\*\*\**P* < 0.0001. (D) Relative survival of *A. baumannii* 17978 wild-type harboring the empty plasmid vector (pVector) or the *pbp1a*-expressing plasmid (pPBP1a) after treatment with 10 mM H_2_O_2_ for 2 h. Data are presented as means ± SD from n= 3. *P* values were determined using an unpaired Student’s t-test. \*\*\*\**P* < 0.0001. (E) Fluorescence (upper) and phase (lower) microscopy of *A. baumannii* 17978 wild-type after treatment with a sublethal dose of sulbactam (0.5x MIC) for 2 h. Scale bar = 2 µm. NADA-green is integrated into the peptidoglycan of living bacteria during cell wall synthesis (45). (F) Surface area quantifications of each cell population (n ≥ 500) were calculated using ImageJ software with the MicrobeJ plugin. Each dot represents one bacterial cell. *P* values were determined using an unpaired Student’s t-test. \*\*\*\**p* < 0.0001. (G) ROS uptake of *A. baumannii* 17978 wild-type after treatment with a sublethal dose of sulbactam (0.5x MIC) for 2 h, followed by 10 mM H_2_O_2_ for 30 min. Data are presented as means ± SD from n = 3. *P* values were determined using an unpaired Student’s t-test. \*\**P* < 0.01. (H) Relative survival of *A. baumannii* 17978 wild-type untreated or treated with a sublethal dose of sulbactam (0.5x MIC), then 10 mM H_2_O_2_ for 2 h. Data are presented as means ± SD from n = 3. P values were determined using an unpaired Student’s t-test. \*\*\**P* < 0.001.

### PBP1a is sufficient to rescue filamentous morphology and bacterial survival by inactivating *iscR*

IscR directly activates *pbp1a* expression (Fig. 2), thereby maintaining coccoid shape under oxidative stress (Fig. 1). PBP1a-mediated coccoid cell shape decreases the uptake of ROS and enhances bacterial resistance to oxidative stress (Fig. 3). Thus, we examined whether PBP1a can restore filamentous to coccoid shape in the *iscR* mutant strain to enhance bacterial survival by decreasing cell surface area during oxidative stress.

Filamentous morphology of *iscR* and *pbp1a* mutants significantly increased in ROS uptake compared to the wild-type and *pbp1b* mutant (Fig. 4A). The increased ROS uptake rendered *iscR* and *pbp1a* mutants more susceptible to oxidative stress (Fig. 1B and S1A). In contrast, inactivation of *pbp1b* did not affect bacterial survival (Fig. S1B). Also, the *pbp1a* mutant with a plasmid expressing *pbp1a* via a heterologous promoter restored bacterial survival under oxidative stress (Fig. S1B). In addition, the *iscR* mutant harboring a plasmid expressing *pbp1a* via a heterologous promoter displayed a shortened cell shape (Fig. 4B) and reduced bacterial surface (Fig. 4C). As expected, the *iscR* mutant with a plasmid expressing *iscR* via a heterologous promoter demonstrates restored cellular morphology and cellular surface (Fig. 4B and C). Furthermore, the decrease in bacterial surface area due to expression of *pbp1a* and *iscR* genes in the *iscR* mutant strain led to reduced ROS uptake (Fig 4D), which subsequently improved bacterial survival under oxidative stress (Fig. 4E). By contrast, the *iscR* mutant with a plasmid expressing *pbp3*, a different peptidoglycan synthase via a heterologous promoter, failed to correct the filamentous shape and bacterial survival observed in the *iscR* mutant (Fig. S2A-B). In summary, PBP1a is sufficient to restore the filamentous cell shape and increased bacterial surface area of the *iscR* mutant strain, thereby decreasing ROS uptake and enhancing bacterial survival.

**Figure 4.**
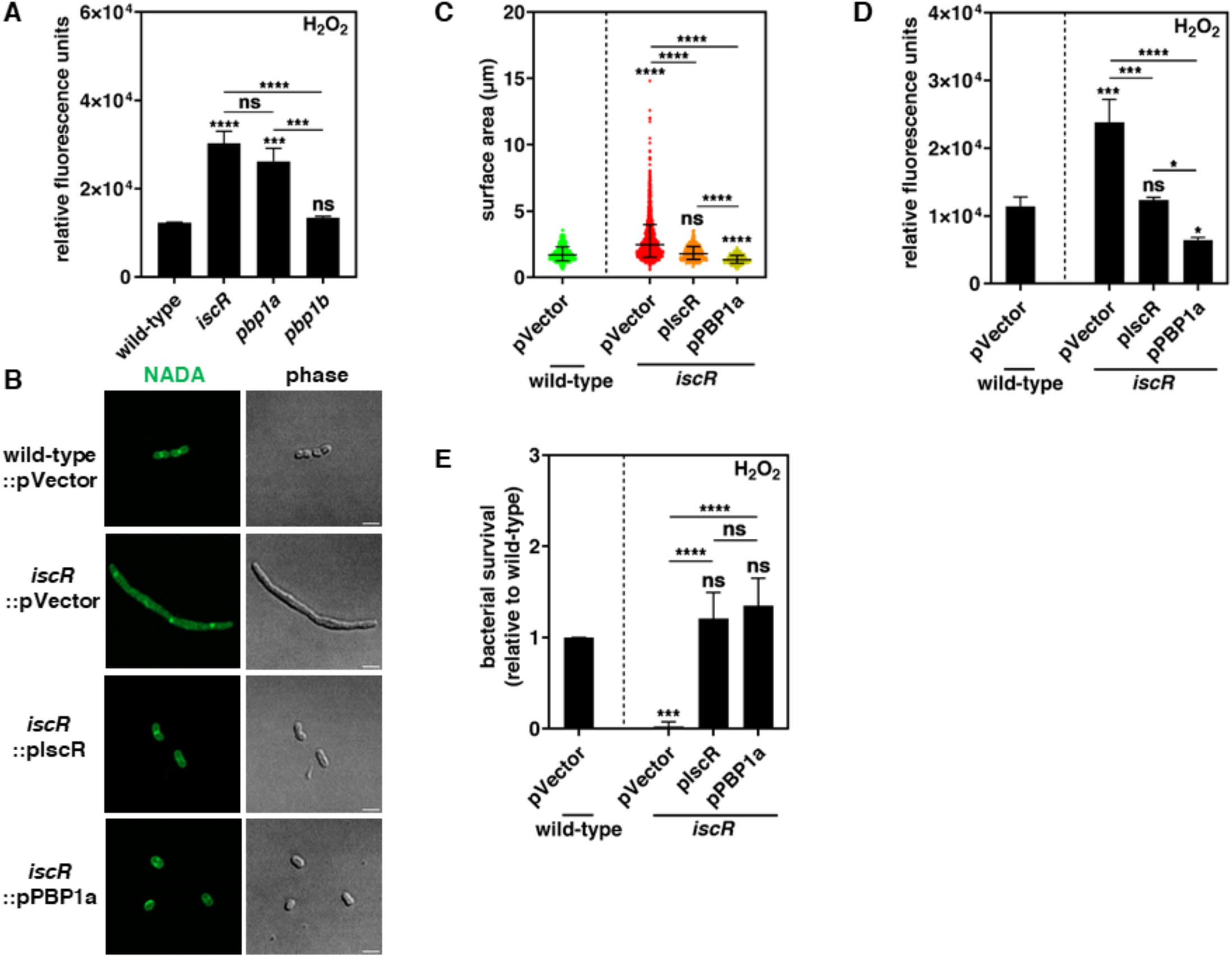
**The *pbp1a* expression rescues the defect in survival of *iscR* mutant under oxidative stress by inducing a change to coccoid shape.** (A) ROS uptake of *A. baumannii* 17978 wild-type, *iscR*, *pbp1a*, and *pbp1b* upon treatment with 10mM H_2_O_2_ for 30 min. The results are expressed as relative fluorescence units, with the RFU value for the untreated control subtracted. Data are presented as means ± SD from n = 3. *P* values were determined using one-way ANOVA with Tukey’s multiple comparison test. \*\*\*\**P* < 0.0001, ns; not significant. (B) Fluorescence (left) and phase (right) microscopy of *A. baumannii* 17978 wild-type harboring the empty plasmid vector (pVector) and *iscR* harboring the empty plasmid vector (pVector), the *iscR*-expressing plasmid (pIscR), or the *pbb1a*-expressing plasmid (pPBP1a) in normal conditions. NADA-green is integrated into the peptidoglycan of living bacteria during cell wall synthesis (45). Scale bar = 2 µm. (C) Surface area quantifications of each cell population (n ≥ 500) were calculated using ImageJ software with the MicrobeJ plugin. Each dot represents one cell. *P* values were determined using one-way ANOVA with Tukey’s multiple comparison test. \*\*\*\**P* < 0.0001, ns; not significant. (D) ROS uptake of *A. baumannii* 17978 wild-type harboring the empty plasmid vector (pVector) and *iscR* harboring the empty plasmid vector (pVector), the *iscR*-expressing plasmid (pIscR), or the *pbb1a*-expressing plasmid (pPBP1a) upon treatment with 10 mM H_2_O_2_ for 30 min. Data are presented as means ± SD from n = 3. *P* values were determined using one-way ANOVA with Tukey’s multiple comparison test. \*\**P* < 0.01, \*\*\**P* < 0.001, ns; not significant. (E) Relative survival of *A. baumannii* 17978 wild-type harboring the empty plasmid vector (pVector) and *iscR* harboring the empty plasmid vector (pVector), the *iscR*-expressing plasmid (pIscR), or the *pbb1a*-expressing plasmid (pPBP1a) after treatment with 10 mM H_2_O_2_ for 2h. Data are presented as means ± SD from n = 3. *P* values were determined using one-way ANOVA with Tukey’s multiple comparison test. \*\*\**P* < 0.001 and \*\*\*\**P* < 0.0001, ns; not significant.

### IscR-mediated *pbp1a* expression reduces the surface area that confers antibiotic resistance

Bacteria can encounter various stresses, including oxidative stress and exposure to antibiotics, during infections. These stress molecules must penetrate the bacterial membrane to inhibit bacterial survival and growth. Most antibiotics target components of the cytoplasm by penetrating the bacterial membrane, except for *β*-lactam antibiotics. Thus, we hypothesized that *A. baumannii* may reduce antibiotic uptake by decreasing cellular surface area via IscR-mediated *pbp1a*expression, as with ROS.

Multiple lines of evidence support the notion that IscR-mediated *pbp1a* expression reduces antibiotic uptake by decreasing cell surface area. First, *A. baumannii* undergoes a transformation to a coccoid shape (Fig. 5A) and reduction of surface area (Fig. 5B) when exposed to tobramycin, an antibiotic that binds to 30S and 50S ribosomal subunits in the cytoplasm (50). Second, the elongated shape of *iscR* allowed quick diffusion of tobramycin into bacterial cytoplasm, as compared to rod-shaped wild-type (Fig. 5C). Third, the reduction in bacterial surface area resulting from PBP1a and IscR expression in the *iscR* mutant strain (Fig. 4B) enhanced bacterial survival against cytosolic antibiotics such as tobramycin, rifampicin, and ciprofloxacin (Fig. 5D-F). In contrast, the *iscR* mutant harboring a plasmid expressing *iscR* or *pbp1a* via a heterologous promoter did not enhance survival when treated with meropenem, a *β*-lactam antibiotic (Fig. 5G). In conclusion, IscR-mediated expression of *pbp1a* is essential for bacterial survival during antibiotic treatment by reducing the cellular surface and maintaining coccoid cellular morphology. This indicates that regulation of cell morphology is critical for defense from various stress molecules, and this is not limited to ROS.

**Figure 5.**
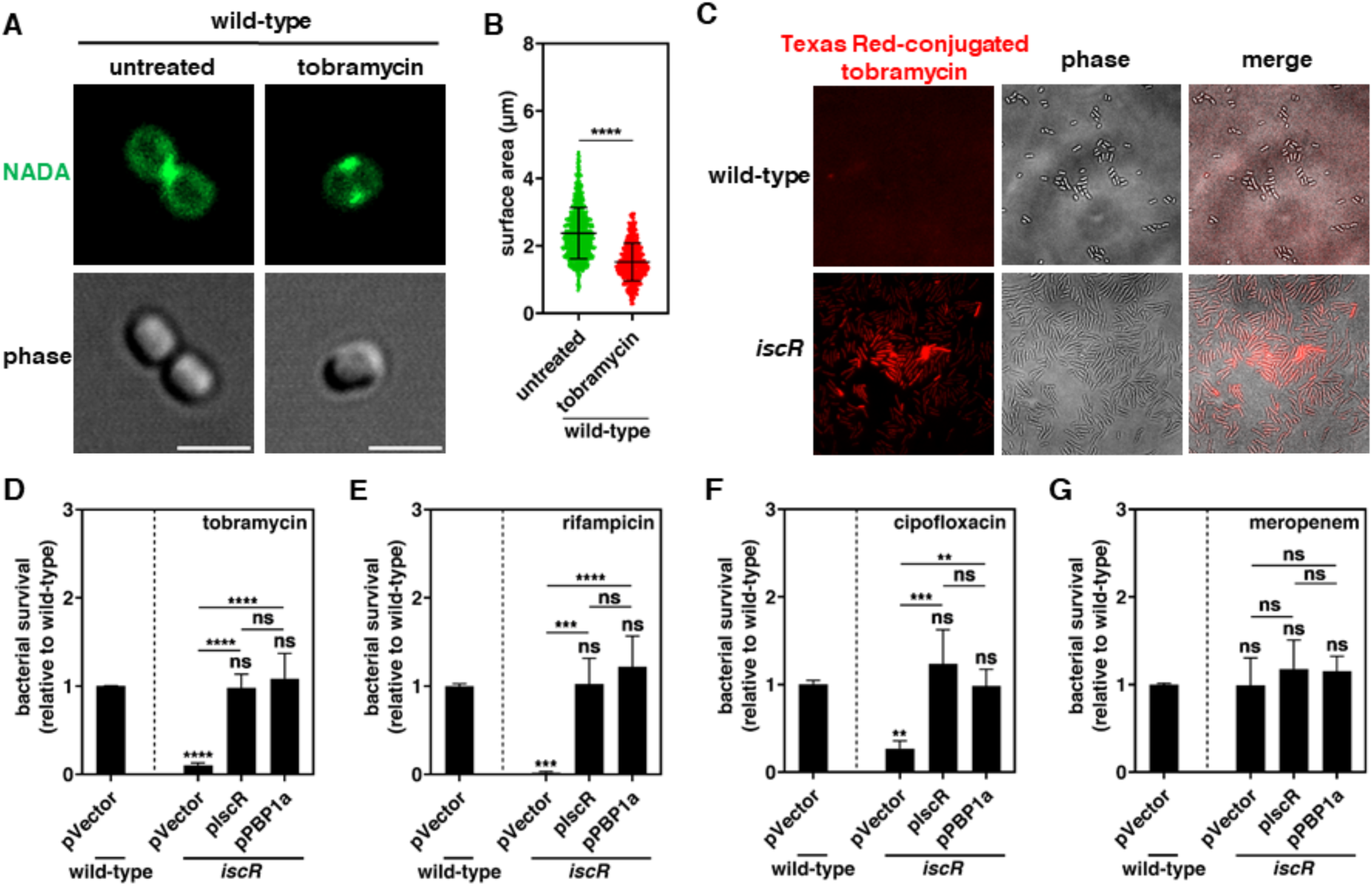
IscR and PBP1a-mediated coccoid shape reduce the uptake of ROS in the cytoplasm in *A. baumannii*. (A) Fluorescence (upper) and phase (lower) microscopy of *A. baumannii* 17978 wild-type untreated or treated with 2x MIC of tobramycin for 30 min. NADA-green is integrated into the peptidoglycan of living bacteria during cell wall synthesis (45). Scale bar = 2 µm. (B) Surface area quantifications of each cell population (n ≥ 500) were calculated using ImageJ software with the MicrobeJ plugin. Each dot represents one cell. *P* values were determined using an unpaired Student’s t-test. \*\*\*\**P* < 0.0001. (C) Uptake of Texas Red-conjugated tobramycin in A. *baumannii* 17978 wild-type and *iscR* at 30 min post incubation with 2x MIC of the conjugated antibiotic. Fluorescence microscopy images are representative of 3 independent experiments. (D-E) Relative survival of *A. baumannii* 17978 wild-type harboring the empty plasmid vector (pVector) and *iscR* harboring the empty plasmid vector (pVector), the *iscR*-expressing plasmid (pIscR) or the *pbb1a*-expressing plasmid (pPBP1a) after treatment with 2x MIC of tobramycin (D), rifampicin (E), ciprofloxacin (F) or meropenem (G) for 2 h. Data are presented as means ± SD from n = 4. *P* values were determined using one-way ANOVA with Tukey’s multiple comparison test. \*\**P* < 0.01, \*\*\**P* < 0.001 and \*\*\*\**P* < 0.0001, ns; not significant.

### IscR-mediated *pbp1a* expression is critical for bacterial survival inside macrophages

Host innate immune cells, including macrophages and neutrophils, kill pathogenic bacteria through oxidative stress, including the production of nitric oxide and hydrogen peroxide (11). We demonstrated that IscR and PBP1a are required for oxidative stress defense by regulating cellular morphology (Figs. 1 and 4). Based on these findings, we reasoned that IscR and PBP1a are essential for bacterial survival within host innate immune cells during infection.

The survival rates of wild-type *A. baumannii* were approximately 3-fold and 5-fold higher than those of *iscR* and *pbp1a*mutants within macrophages, respectively, at 6 hours post-phagocytosis (Fig. 6A). In addition, bacterial survival was significantly enhanced in *iscR* or *pbp1a* mutants harboring plasmids via expressing their respective genes from heterologous promoters (Fig. 6B). By contrast, isogenic strains with empty plasmid did not restore bacterial survival inside macrophages (Fig. 6B). Consistent with *pbp1a* overexpression can restore bacterial morphology and enhance survival of the *iscR* mutant under oxidative stress conditions (Fig. 4), expression of *pbp1a* via heterologous promoters in the *iscR* mutant partially restored bacterial survival within macrophages (Fig. 6B). Because IscR is essential for bacterial survival by regulating genes necessary for adapting to multiple host-imposed stresses such as iron limitation and oxidative stress in *Vibrio vulnificus* (36) and *Salmonella enterica* (51), PBP1a expression partially rescue bacterial survival defect of *iscR* mutant strain inside macrophage during infections. Additionally, the inactivation of *iscR* reduced bacterial survival in spermine, a recognized generator of the oxidative stress molecule nitric oxide, compared to the wild-type (Fig. 6C). The survival defect induced by spermine in the *iscR* mutant was corrected by a plasmid expressing the *iscR* gene, whereas the empty plasmid did not rescue bacterial survival (Fig. 6D). Altogether, IscR-mediated expression of *pbp1a* is essential for bacterial survival within host innate immune cells during infections.

**Figure 6.**
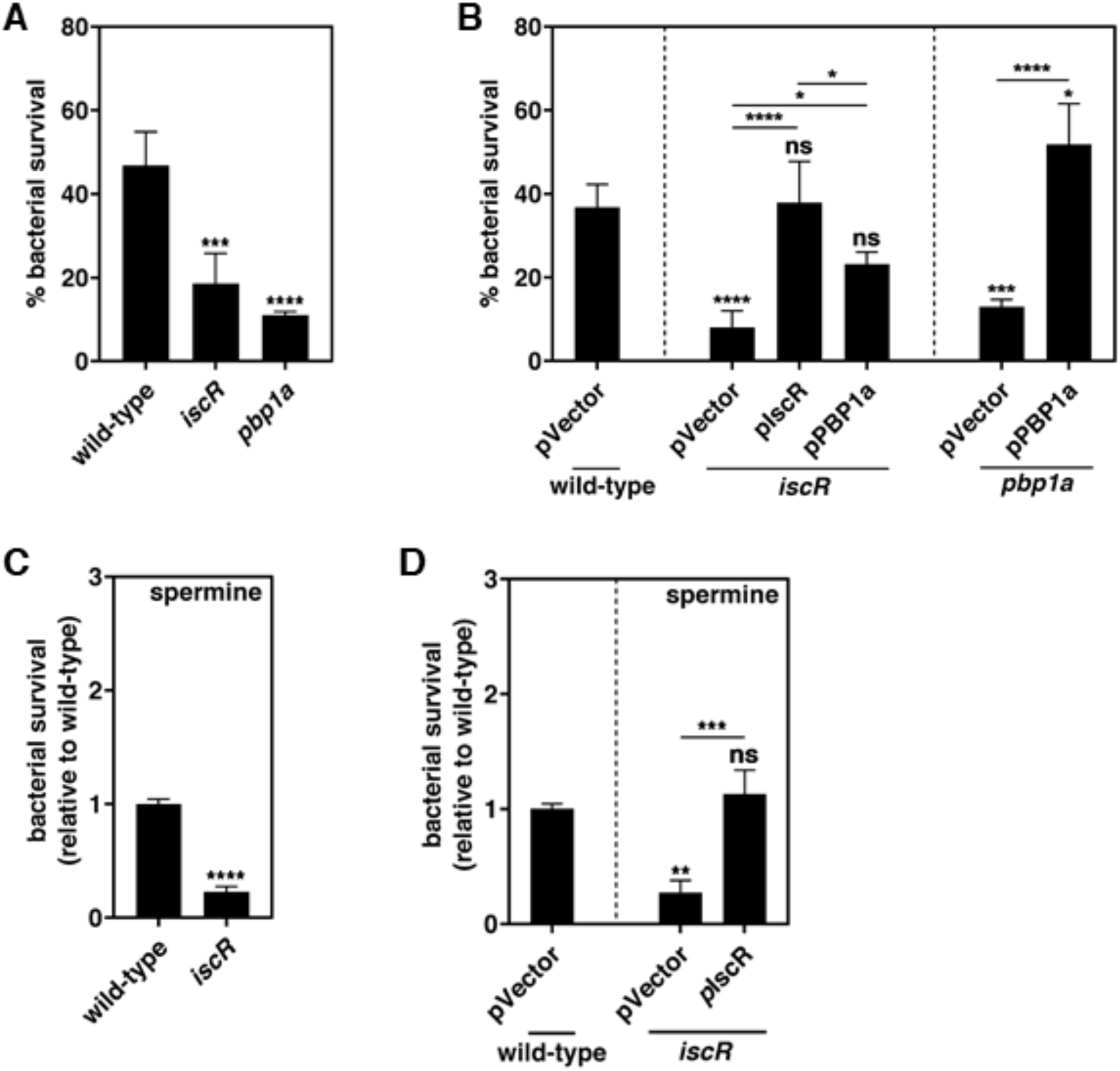
Both IscR and PBP1a are required for bacterial survival inside macrophages. (A) Survival of *A. baumannii* 17918 wild-type, *iscR* mutant, and *pbp1a* mutant inside J774A.1 macrophage at 6 h post-infection. (B) Survival of *A. baumannii* 17918 wild-type harboring the empty plasmid vector (pVector), *iscR* mutant harboring the empty plasmid vector (pVector), the *iscR*-expressing plasmid (pIscR) or *pbp1a*-expressing plasmid (pPBP1a), and *pbp1a* mutant harboring the plasmid vector (pWH1266) or the *pbp1a*-expressing plasmid (pPBP1a) inside J774A.1 macrophage at 6 h post-infection. (C) Relative survival of *A. baumannii* 17978 wild-type and *iscR* after treatment with 1 mM spermine NONOate. (D) Relative survival of *A. baumannii* 17978 wild-type harboring the empty plasmid vector (pVector) and *iscR* mutant harboring the empty plasmid vector (pVector) or the *iscR*-expressing plasmid (pIscR) after treatment with 1 mM spermine NONOate. Data are presented as means ± SD from n = 4 (A and B) or n = 3 (C and D). *P* values were determined using one-way ANOVA with Dunnett’s multiple comparison test (A), or one-way ANOVA with Tukey’s multiple comparison test (B and D), or unpaired Student’s t-test (C). *P < 0.05, **P < 0.01, ***P < 0.001 and ****P < 0.0001, ns; not significant.

## DISCUSSION

In this study, we demonstrate that the transcriptional regulator IscR protects *Acinetobacter baumannii* against oxidative stress (Fig. 1) and antibiotic treatment (Fig. 5) by regulating cellular morphology. IscR directly activates the expression of *pbp1a*, a peptidoglycan synthase (Fig. 2) that facilitates the transition of cell shape from elongated rod shape to compacted coccoid shape in response to reactive oxygen species (ROS) and antibiotics (Fig. 2-4). This morphological change reduces bacterial surface area, thereby restricting the uptake of stress molecules and increasing bacterial survival under stress conditions (Figs. 4 and 5). By contrast, inactivation of *iscR* or *pbp1a* displayed filamentous morphology that enhanced stress-molecule uptake (Fig. 4). Notably, IscR-mediated morphology regulation is critical for pathogenesis inside macrophages (Fig. 6) and murine model (Fig. 1D). Therefore, IscR-mediated remodeling of bacterial shape is essential for bacteria to resist against immune defenses and antibiotic treatment during infection (Fig. 7).

**Figure 7.**
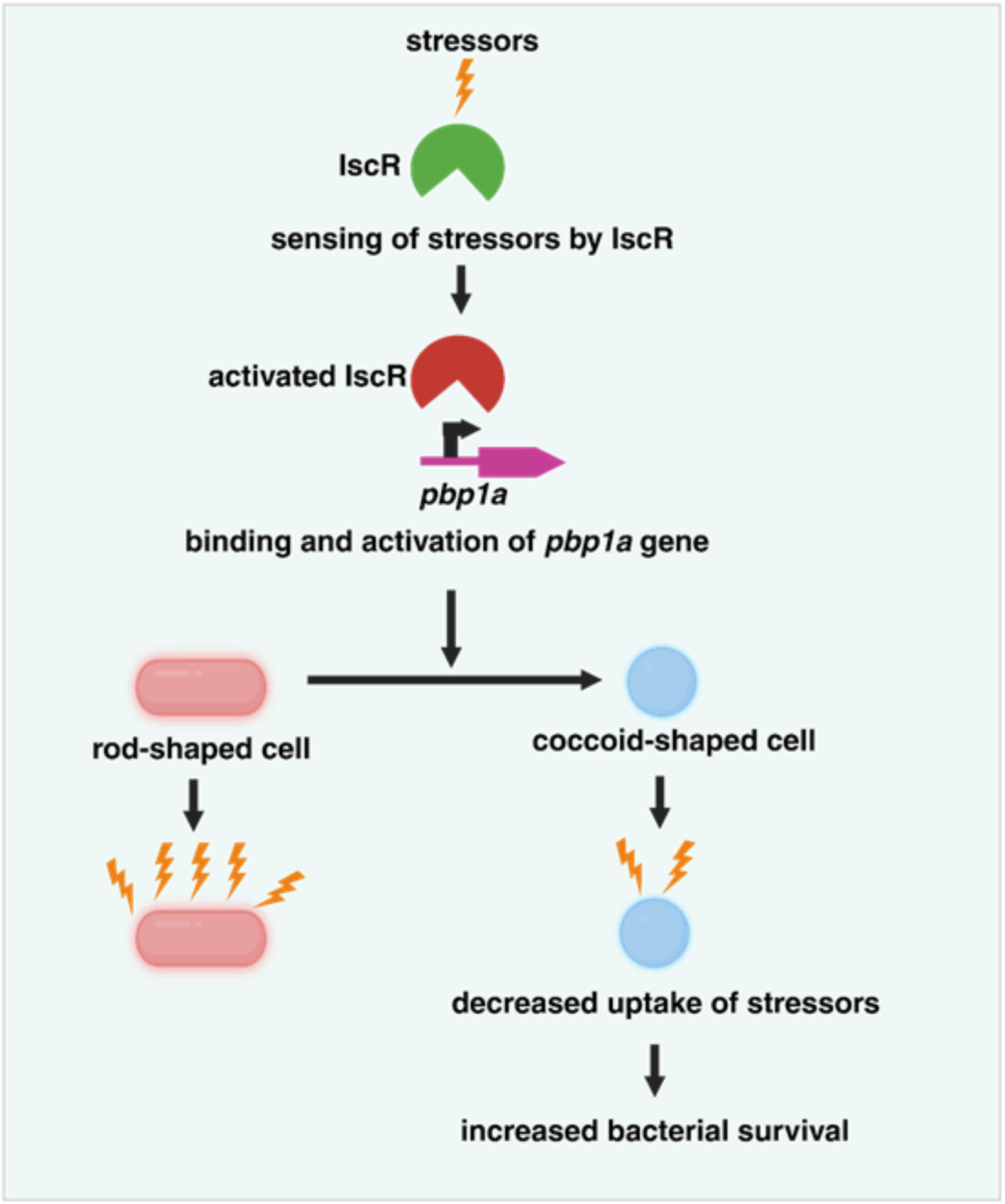
IscR-mediated morphological regulation confers stress resistance by reducing the uptake of stress molecules in *A. baumannii*. IscR is activated by oxidative stress or antibiotics. Then, IscR activates *pbp1a* expression, causing the transition from rod shape to coccoid shape. The coccoid shape allows bacteria to reduce the entry of stress molecules into the cytoplasm, thereby increasing bacterial survival within the host during infection and antibiotic treatment.

All three domains of life regulate cellular shape to adapt to environmental conditions. Although a decreased surface area, such as in a coccoid or round cell shape, generally reduces the efficiency of transporting materials between environments (52, 53), it also offers multiple advantages for organisms. In animals, a lower surface area is better at retaining heat, minimizing heat loss to the environment (54, 55). For instance, animals continuously exposed to cold temperatures decrease their surface area to reduce heat loss and maintain body temperature (54, 55). In addition, plants try to reduce their surface area to help prevent desiccation in dry periods (56, 57). Plants and insects reduce surface area to avoid water loss during osmotic stress (58, 59). Moreover, a reduced surface area can easily store more nutrients or energy than a large-surface-area cell shape under nutrient-limited conditions (3, 60), because a smaller surface area can protect its internal resources from external fluctuations. Therefore, surface reduction in unfavored conditions is a widely conserved defense mechanism in all living organisms. Here, we investigated that bacteria reduce their surface area to inhibit the uptake of stress molecules, thereby protecting themselves from environmental stress (Fig. 7).

In many bacterial species, rod-shaped cells change to a coccoid shape under stress conditions, including nutrient starvation, low pH, and the stationary phase (1–5). Coccoid cell formation in *Pseudomonas aeruginosa* is induced by carbapenem and penicillin antibiotics (61). Uropathogenic *Escherichia coli* (UPEC) also undergoes morphological changes from a rod to a coccoid shape inside the autophagosome of human bladder epithelial cells (62). Thus, the morphological transition from rod to coccoid shape is regarded as a bacterial survival strategy in response to stress. However, it is poorly understood how the coccoid shape contributes to the survival and pathogenesis of pathogenic bacteria during infection. This study demonstrates that viable coccoid cells reduce the uptake of stress molecules, such as ROS and antibiotics, by decreasing bacterial surface area through IscR-mediated PBP1a production in the pathogenic bacterium *Acinetobacter baumannii* (Fig. 2-4). Furthermore, this is required for survival and pathogenesis during infection in host innate immune cells (Fig. 6) and in the mouse model (Fig 1D). Our findings suggest that the transition to a viable coccoid cell shape under unfavorable conditions may hinder the entry of stress molecules into the bacterial cytoplasm, a common strategy for the survival of pathogenic bacteria during infection (Fig. 7).

The emergence of multidrug-resistant bacteria poses a severe threat to public health, requiring the development of innovative strategies for antimicrobial therapy. Our findings indicate that drug combinations with conventional antibiotics and inhibitors for bacterial cell shape regulation may effectively eliminate bacteria. In a clinical setting, combination therapy with the *β*-lactamase inhibitor sulbactam and the *β*-lactam antibiotic ampicillin was widely used for decades to treat bacterial infections (63). Our study revealed that bacteria fail to transition to the coccoid form and remain filamentous under sulbactam treatment (Fig. 3E) and filamentous forms easily uptake antibiotics (Fig. 4). Thus, they are more valuable than coccoid cells under a *β*-lactam antibiotic treatment (Fig. 7). Consequently, the combination of sulbactam with other antibiotics is expected to demonstrate strong bactericidal effects against bacteria at concentrations where each individual drug shows minimal bactericidal activity. Notably, recent studies indicate that sulbactam combinations with various antibiotics, including colistin and durlobactam, enhance treatment outcomes for carbapenem-resistant *A. baumannii* (CRAB) (64–67), a pathogen the WHO has designated as the most critical (68). Therefore, our findings establish a framework for identifying novel antibacterial drug combinations to treat antibiotic-resistant bacterial infections by promoting the uptake of conventional antibiotics by pathogens.

## MATERIALS AND METHODS

### Bacterial strains and growth conditions

Bacterial strains and plasmids employed in this study are listed in Table S1. For routine cultivation and genetic manipulation, *A. baumannii* and *E. coli* were grown at 37°C in Luria Bertani (LB) broth or on LB agar (BD Difco). For selection of recombinants or maintenance of plasmids in *A. baumannii* and *E. coli*, antibiotics were supplied at the following concentrations: ampicillin, 50 µ g/mL; 100 carbenicillin, 100 µ g/mL; gentamicin 10 µg/mL; and kanamycin, 50 µg/mL.

### Cell lines

The J774A.1 mouse macrophage cell line was cultured in Dulbecco’s Modified Eagle Medium (DMEM) High Glucose (Welgene) supplemented with 10% heat-inactivated fetal bovine serum (Welgene) at 37°C and 5% CO_2_.

### Mouse models

All animal studies were approved from the Institutional Animal Care and Use Committee (IACUC) at the Seoul National University. Wild-type C57BL/6 mice (Orient Bio, Seongnam, South Korea) were housed and maintained in the specific pathogen-free facility at Seoul National University College of Medicine.

### Construction of chromosomal mutants and plasmids in *A. baumannii*

Primers used in this study are listed in Table S2. Chromosomal mutants were generated using standard protocols described previously (69). Briefly, the FRT-flanked kanamycin resistance determinant of pKD4 was amplified using primers bearing ∼50 bp of homology to the region flanking the gene of interest. This PCR product was electroporated into competent *A. baumannii* carrying pAT02 plasmid, which expresses the RecAb recombinase. Mutants were selected on LB agar plates with kanamycin, and integration of the resistance marker was confirmed by PCR. To remove kanamycin resistance cassette, electrocompetent mutants were transformed with pAT03 plasmid, which expresses the FLP recombinase. All mutant strains were confirmed by PCR and sequencing.

Complementation of *iscR* or *pbp1a* mutants was performed *in trans* using the vector pWH1266 (70). The gene and its native promoter were amplified using genomic DNA from wild-type ATCC 17978 as template and the primers listed in table S2. pWH1266 was digested with BamHI and SalI (New England Biolabs). The purified PCR product and cut plasmid were joined using NEB Builder Hifi Assembly (New England Biolabs) at 50°C for 15 min. The resulting assembly was transformed into chemically compenent DH5α by heat shock, and successful transformants were selected on LB agar plates with ampicillin following overnight growth at 37°C. The constructs were confirmed by PCR and sequencing prior to introduction of this vector, as well as the native empty vector, to the mutants. Successful transformants were selected on LB agar plates with ampicillin.

### Bacterial survival assays

Freshly streaked bacterial strains were grown overnight in LB broth at 37°C under shaking. Overnight cultures were sub-cultured at an OD_600_ of 0.005 in LB broth and grown to early-stationary phase (6 h). Bacteria were diluted to an OD_600_ of 0.5, washed twice with 1X phosphate buffered saline (PBS, pH 7.4) (137 mM NaCl, 2.7 mM KCl, 10 mM Na_2_HPO_4_, and 1.8 mM KH_2_PO_4_). Then bacteria were re-suspended in PBS without or with drugs and incubated at 37°C with shaking for 2 h. The drugs include H_2_O_2_, and Spermine NONOate (GlpBio). Name of the drug and its concentration are indicated in figures. The live bacterial numbers or colony-forming units (CFUs) were determined by serial dilution and plating. Bacterial survival was calculated by dividing the number of CFU of drug-treated bacteria by the number of CFU of non-treated bacteria.

### Fluorescent NADA staining and image analysis

To visualize bacterial peptidoglycan, a previously described protocol has been used (26, 46, 71). Briefly, sub-cultured bacteria were washed once with LB broth, incubated with 3 µL of 10 mM NBD-(linezolid-7-nitrobenz-2-oxa-1,3-diazol-4-yl)-amino-D-alanine (NADA) (Tocris Bioscience) at 37°C for 30 min, and fixed with 1X PBS containing 4% paraformaldehyde (Biosesang). Fixed bacteria were immobilized on agarose pads and imaged using Zeiss LSM 880 inverted confocal microscope or Zeiss AxioObserver.Z1 inverted fluorescence microscope with a 63X oil immersion objective and a laser line of 488 nm for detection. All images were processed using ImageJ Fiji (72), and the MicrobeJ plug-in was used for quantifications of bacterial surface (73). At least 300 cells were analyzed for all experiments and data were plotted using GraphPad Prism 10 (10.4.1). Each experiment was repeated three times, and each replication was used in the data set. One representative image from the data sets was included in each figure.

### Reporter assays

To construct reporter plasmids (pLPV1Z-P*_pbp1a_* and pLPV1Z-P*_isc operon_*), promoter regions of the *pbp1a* operon and the *isc* operon, respectively, were amplified using the primers listed in Table S2 (74). pLPV1Z was digested with PstI and XbaI (New England Biolabs). The purified PCR product and cut plasmid were joined using NEB Builder Hifi Assembly (New England Biolabs) at 50°C for 15 min. The resulting assembly was transformed into chemically compenent DH5α by heat shock, and successful transformants were selected on LB agar plates with gentamicin following overnight growth at 37°C. The constructs were confirmed by PCR and sequencing prior to introduction of this vector to wild-type and mutants. Successful transformants were selected on LB agar plates with gentamicin. For experiments to check gene expression (Fig. 2D and E), bacterial strains harboring pLPV1Z-P*_pbp1a_* or pLPV1Z-P*_isc operon_* were grown overnight in LB broth at 37°C under shaking. Overnight cultures were sub-cultured at an OD_600_ of 0.005 in LB broth and grown to early-stationary phase (6 h). 100 µL of bacterial culture was added to 100 µL of LB broth without and with 10 mM of H_2_O_2_ in 96-well white plate. The luminescence and OD_600_ value were recorded using the BioTek Synergy H1 plate reader.

### IscR protein purification

Full-length IscR of *A. baumannii* was obtained though PCR and inserted into the pET28b (+) vector with 6x His tag at the N-terminus. pET28B-PmrA was transformed into BL21 (DE3) *E. coli* competent cells. The cells were grown overnight in LB broth at 37°C with shaking. Overnight culture was sub-cultured 1:100 in LB broth and grown at 37°C with shaking to an OD_600_ of 0.4. Culture was then induced with 1 mM of isopropyl β–D-1-thiogalactopyranoside (IPTG, Sigma-Aldrich) for 1 h at 30°C with shaking. Cells were harvested and pellet was resuspended in binding buffer (pH 7.4) containing tris buffered saline (TBS, Biosesang) and 10 mM of Imidazole (Sigma-Aldrich). Cells were lysed by sonication, and the resulting clarified supernatant was loaded onto a Econo-Pac Chromatography Column (Bio-Rad) which has been pre-loaded with Nickel Sepharose High Performance (Cytiva) and equilibrated with binding buffer. After overnight incubation at 4°C with rotating, the column was washed three times with wash buffer (pH 7.4) containing TBS and 40 mM Imidazole. IscR protein was eluted using elution buffer (pH 7.4) containing TBS and 250 mM Imidazole. Fractions containing pure protein were then collected after SDS-PAGE, combined and concentrated using Amicon Ultra centrifugal filters (Merck Millipore). Protein concentration was determined using BCA Protein Assay kit (Pierce) according to the manufacturer’s protocol.

### Electrophoretic mobility shift assay (EMSA)

EMSA assays were performed using the LightShift Chemiluminescent EMSA kit (Thermo Scientific). Biotinylated DNA fragments from the upstream regions of *pbp1a, isc* operon, and *hscB-hscA-fdx* operon were obtained by PCR using 5’-end biotin-labeled primers listed in Table S2. Binding reactions were incubated in 20 µL volume containing 25 fmol of biotinylated DNA and various concentrations of purified IscR protein (0 – 1.5 µM) in 1x Gel Shift Binding Buffer (Molecular Depot) for 60 min. To ensure binding specificity, a 50-fold molar excess of unlabeled competitor DNA was added to the reaction mixture where indicated. Binding reactions were loaded with the provided loading dye onto a 6% Novex TBE Gel (Invitrogen) and separated in 0.5x Novex TBE Running Buffer (Thermo Fisher) at 100 V for 60 min at 4°C. A Trans-blot Turbo Transfer System (Bio-Rad) was used to transfer proteins and DNA to Biodyne B positively charged nylon membranes (Thermo Fisher) for 30 min at 25 V. Cross-liking was performed by exposing the membranes to UV light for 15 min in a Dual LED Blue/White Light Transilluminator (Invitrogen). Blocking, incubation with streptavidin-horseradish peroxidase conjugate, washing and revealing were performed according to the protocol. Membranes were imaged using Amersham ImageQuant 900 (Cytiva).

### ROS uptake assays

2’,7’-dichlorodihydrofluorescein diacetate (DCFH-DA) was utilized as a fluorescent dye to detect ROS production upon treatment of bacteria with H_2_O_2_. Sub-cultured bacteria were washed with 1X PBS and treated with 10 µM of DCFH-DA for 30 min at 37°C. Bacteria then were washed three times with 1X PBS to ensure the removal of excess fluorescent probes. Next, the probed-bacteria were treated with 10 mM H_2_O_2_ for 30 min. After incubation, the fluorescence intensity was measured with excitation and emission wavelengths at 488 and 525 nm.

### Texas Red-conjugated tobramycin uptake assays

To make Texas Red-conjugated tobramycin (TRT), Texas Red™ Sulfonyl Chloride (Invitrogen) was dissolved in anhydrous N, N-Dimethylformamide (DMF) to 5 mg/mL on ice. Tobramycin (Sigma, 10 mg/mL) was mixed with Texas Red at a 1:1 volume ratio (100 µL each) in 0.1 M sodium bicarbonate buffer (800 µL, pH 8.5), followed by rotation and incubation on ice for 2 hours in the dark. The reaction mixture was desalted to remove free dye, and the conjugate was diluted to 1.0 µg/mL for experimental use.

To investigate the time scale of antibiotic accumulation in living bacteria, sub-cultured bacteria were incubated with 1X MIC of TRT for 30 min, immobilized on agarose pads and imaged using Zeiss AxioObserver.Z1 inverted fluorescence microscope with a 63X oil immersion objective and a laser line of 555 nm for detection. All images were processed using ImageJ Fiji. One representative image from three independent experiments was shown.

### Intramacrophage survival Assay

For the intracellular survival assays, J774A.1 cells were seeded in 24-well plates and incubated at 37°C and 5% CO_2_ for 36 h before the experiment (2×10^4^ cells/well). *A. baumannii* strains were grown overnight in LB at 37°C under shaking condition. Overnight cultures were sub-cultured at an OD_600_ of 0.005 in LB and grown to early-stationary phase (6 h). An appropriate volume was added to each well to reach an MOI of 100. Plates were centrifuged 10 min at 700 × g to enhance bacterial contact with host cells and incubated for 1 h at 37°C and 5% CO2. Cells were washed twice with PBS and subsequently incubated with medium supplemented with 100 µg/mL gentamicin to kill extracellular bacteria. At 2 h post infection (p.i.), the concentration of gentamicin was decreased to 10 µg/mL. At 2 h and 6 h p.i., cells were washed twice with PBS and lysed with 500 µL PBS containing 0.2% Triton X-100. Lysates were serially diluted and plated on LB agar plates to determine CFU. Percentage of bacterial survival was calculated by dividing the number of CFU at 6 h p.i. by the number of CFU at 2 h p.i.

### Intraperitoneal infection of A. baumannii strains in mice

For i.p. inoculation in mice, freshly cultured inocula were prepared as previously described. Mice were inoculated with 5×10^8^ CFU/mL of A. baumannii in 200 µL PBS using 1.0 mL BD Ultra-Fine Insulin Syringes with a 31 G needle. Groups of six infected mice were monitored for their survival post inoculation.

### Quantification and statistical analysis

Data and statistical analysis were performed using GraphPad Prism 10.4.1. Replicates and statistical details can also be found in the methods and figure legends.

## ACKNOWLEDGMENTS

We are also grateful to Prof. Bryan Davies at the University of Texas at Austin for providing pAT02 and pAT03 plasmids, and Prof. Paolo Visca at the Roma Tre University for providing pLPV1Z plasmid. This work was supported by the New Faculty Startup Fund from Seoul National University to J.Y., the National Research Foundation Korea Basic Science Research Programs (2021R1C1C1005184, RS-2026-25489275 and 2020R1A5A1019023 to J.Y.), the Global-LAMP Program of the National Research Foundation of Korea (NRF) grant funded by the Ministry of Education (RS-2023-00301976), the Korea Health Technology R&D Project through the Korea Health Industry Development Institute (KHIDI) (HI23C026400 and RS-2023-00304637), and Brain Pool program by the National Research Foundation of Korea (2021H1D3A2A02083176 to H.V.N).

